# Low effective mechanical advantage of giraffes’ limbs during walking reveals trade-off between limb length and locomotor performance

**DOI:** 10.1101/2021.04.29.441773

**Authors:** Christopher Basu, John R. Hutchinson

## Abstract

Giraffes (*Giraffa camelopardalis*) possess specialised locomotor morphology, namely elongate and gracile distal limbs. Whilst this contributes to their overall height (and enhanced feeding behaviour), we propose that the combination of long limb segments and modest muscle lever arms results in low effective mechanical advantage (EMA, the ratio of in-lever to out-lever moment arms), when compared with other cursorial mammals. To test this, we used a combination of experimentally measured kinematics and ground rection forces (GRFs), musculoskeletal modelling, and inverse dynamics to calculate giraffe forelimb EMA during walking. Giraffes walk with an EMA of 0.34 (±0.05 S.D.), with no evident association with speed within their walking gait. Giraffe EMA was markedly below the expectations extrapolated from other mammals ranging from 0.03 – 297 kg, and provides further evidence that EMA plateaus or even diminishes in mammals exceeding horse size. We further tested the idea that limb segment length is a factor which determines EMA, by modelling the GRF and muscle moment arms in the extinct giraffid *Sivatherium giganteum* and the other extant giraffid *Okapia johnstoni. Giraffa* and *Okapia* shared similar EMA, despite a 4-6 fold difference in body mass (*Okapia* EMA = 0.38). In contrast *Sivatherium*, sharing a similar body mass to *Giraffa*, had greater EMA (0.59), which we propose reflects behavioural differences, such athletic performance. Our modelling approach suggests that limb length is a determinant of GRF moment arm magnitude, and that unless muscle moment arms scale isometrically with limb length, tall mammals are prone to low EMA.

**Significance Statement:** Giraffes are the tallest living animals - using their height to access food unavailable to their competitors. It is not clear how their specialized anatomy impacts their athletic ability. We made musculoskeletal models of the forelimbs from a giraffe and two close relatives, and used motion-capture and forceplate data to measure how efficient they are when walking in a straight line. A horse for example, uses just 1 unit of muscle force to oppose 1 unit of force on the ground. Giraffe limbs however are comparatively disadvantaged – their muscles must develop 3 units of force to oppose 1 unit of force at the ground. This explains why giraffes walk and run at relatively slow speeds.

## Introduction

Giraffes (*Giraffa camelopardalis*, Linnaeus 1758) are feeding specialists, but does the possession of a disproportionately long neck and long limbs facilitate or constrain other behaviors? Whilst their anatomy confers a recognized feeding advantage (Cameron and Toit, 2007), the effect on locomotor performance remains unclear. Giraffes embody the essence of cursorial morphology. Cursoriality refers to a number of anatomical traits which lend themselves to enhanced locomotor performance, including elongate distal limbs, digit loss or reduction, and restriction of joint rotation to the parasagittal plane (Gregory, 1912, Coombs, 1978). One method of measuring the degree of cursoriality is the ratio of metatarsal to femur length (MT:F). By this measure, giraffes display extreme cursoriality, with MT:F 1.4 (Garland and Janis 1993). Considering that horses (*Equus ferus caballus*), fast-running and quintessential cursorial mammals, have a MT:F of 0.8, giraffe morphology is extreme.

Mitchell suggested that giraffes’ elongated appendicular skeleton delivers a ‘mechanical advantage’ during locomotion (1), and Pincher speculated that long limbs facilitate fast running speed (2). Yet despite their extreme cursorial morphology, giraffes are athletically challenged. For example adults giraffes run and walk at modest speeds, and lack an aerial phase in their galloping gait (3, 4), conforming to the observation that the largest terrestrial animals are not the fastest (5-7).

We propose that maximal locomotor performance in giraffes is constrained by their elongate limb segments (and consequently high shoulder height), rather than enhanced by it. At increasing distances from the ground, ground reaction force (GRF) vectors are more horizontally distant from the foot’s center of pressure (COP); or point of GRF application. As a result, limb joints in taller animals may be subject to larger GRF moment arms than the homologous joints in shorter animals. Large GRF moment arms may reduce the effective mechanical advantage of the limb, or put more simply, limit the ability to resist gravitational forces (Biewener, 1989,1990). Giraffids (Figure 1A and 1B) are an ideal group in which to explore this idea, as a diverse range of phenotypes (with respect to height) have existed in the lineage.

Effective mechanical advantage (EMA) is a measure of a given joint’s (or limb’s) leverage against the GRF; or in a simpler sense, the relative suitability of the joint (or overall limb) to resist gravity (Figure 2A). EMA is a useful variable to consider in the context of locomotion, as it is inversely proportional to the muscle force required to balance GRFs during locomotion, and is also associated with mechanical stress (8) and activated muscle volumes (9-11). EMA can be expressed as the ratio of the “antigravity” (typically extensor, or joint-straightening) in-lever muscle moment arm (r), to the out-lever moment arm of the GRF vector (R) during the stance phase of locomotion:

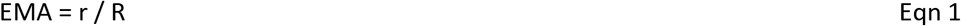

**Figure 1.**
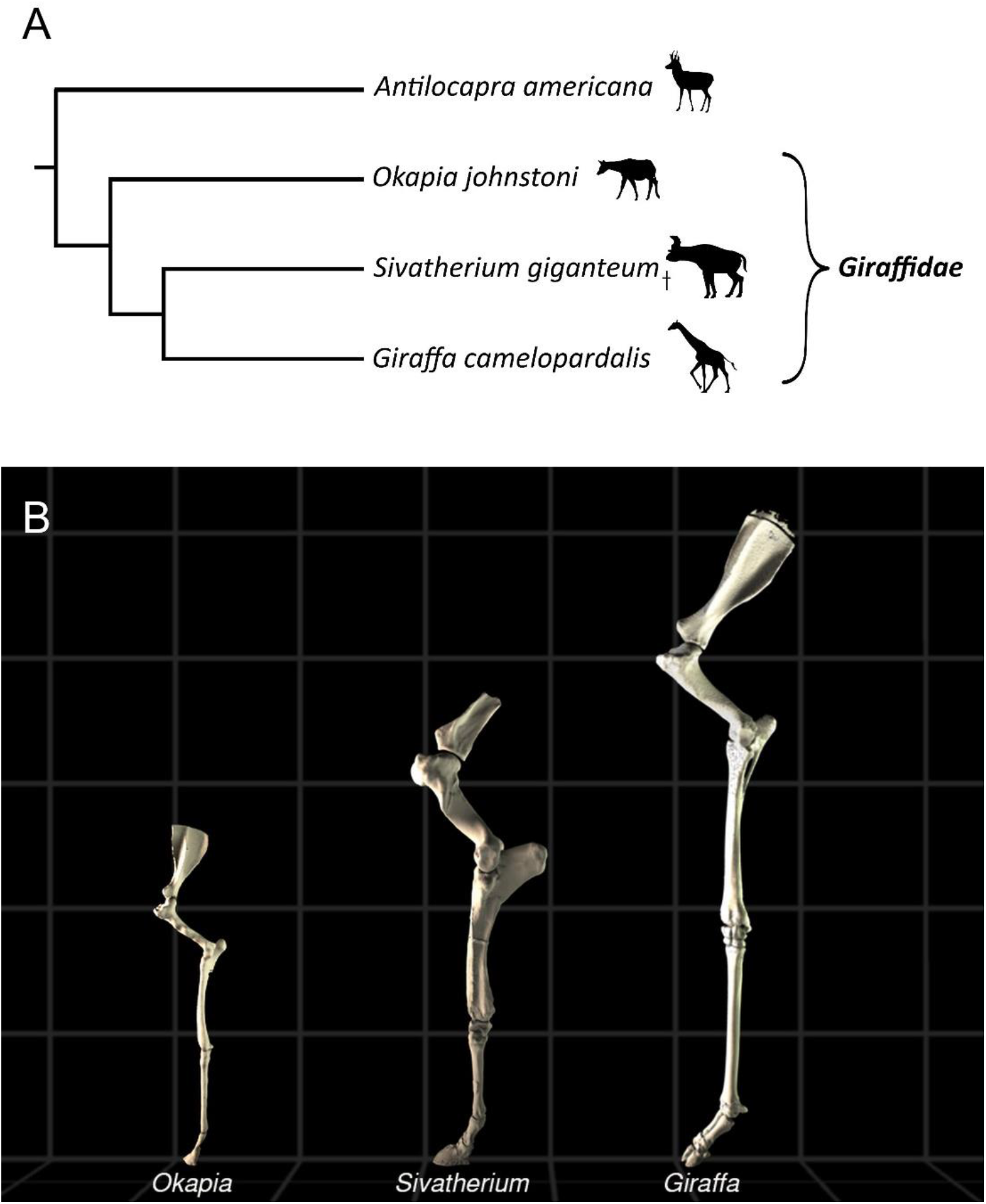
(A) Phylogeny of Giraffidae and outgroup (Ríos, Sánchez et al. 2016); † refers to an extinct taxon. Image credits: https://www.phlopic.org (B) Modelled midstance postures of left forelimbs of *Okapia*, *Sivatherium* and *Giraffa*. Models are displayed to scale, with each gray box measuring 0.5 m in length.

**Figure 2.**
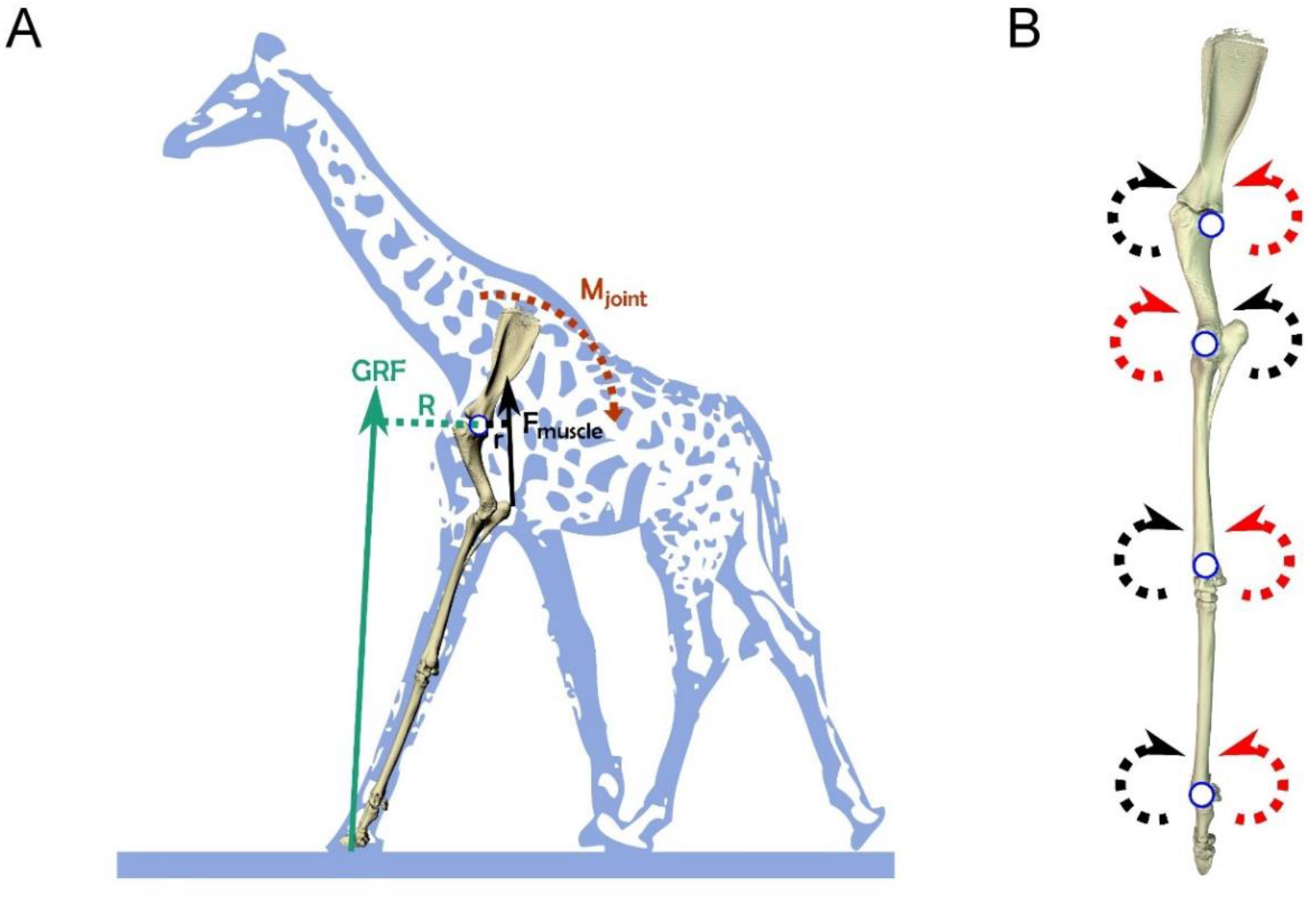
(A) Giraffe forelimb skeleton during the early stance phase, with associated GRF vector (green arrow), originating from a point (COP) under the foot. The GRF vector has a moment arm (R; green dotted line) with respect to the shoulder joint, inducing a joint moment (M_joint_). To resist this, muscle force (F_muscle_) produces an opposing muscle moment, with moment arm r (short black dotted line). (B) Locations of joint centers used to set up a coordinate system for the giraffe musculoskeletal model (left forelimb in lateral view). Red arrows represent flexion; black arrows represent extension.

EMA scales allometrically with body mass in mammals ranging from mice (0.03 kg) to horses (275 kg), with a scaling exponent of 0.26 (12). This indicates that larger animals exert relatively smaller muscle forces in order to resist gravitational collapse of their limbs during the stance phase (here, with EMA measured at the trot-gallop transition). Horses have an EMA of approximately 1, indicating that their extensor muscle moment arms are equal to their GRF moment arms, on average. Hence for every 1 N of GRF, horses typically must develop 1 N of muscle force to maintain their posture. Their large EMA can be explained by their relatively upright posture, where their joints are closely aligned with the GRF vector.

A plateau might exist in the relationship of EMA with body mass, in animals exceeding horse size. Asian elephants (*Elephas maximus*) have an EMA of approximately 0.68 during slow walking (11), and a musculoskeletal model of the extinct *Tyrannosaurus rex* estimated that this animal moved with similar EMA (13). Similarly, relatively straight-limbed humans walk with an EMA ~0.7 (Biewener et al., 2004); and both humans and elephants shift to EMA ~0.5 or less during more crouched running gaits (Ren, Miller et al. 2010). Hence horses have the highest EMA yet recorded, partly explaining their high athletic capacity despite their large size (e.g., Garland, 1983).

The evolution of the giraffid appendicular skeleton has functional implications involving EMA. Giraffids with more ancestral morphology (Figure 1A) possessed relatively shorter limb segments, and smaller body mass than *Giraffa* (14, 15). Okapis (*Okapia johnstoni*), the only other living giraffids, have body proportions considered to be more ancestral; with a modest body mass of 250 kg (16), and moderate limb and neck elongation (14, 17, 18).

*Sivatherium giganteum* (Falconer and Cautley 1836), from an extinct giraffid lineage (Figure 1A), displayed a different morphological phenotype, featuring extreme body mass in the presence of a robust appendicular skeleton and short neck (19). Comparing the EMA of giraffes, okapis and *Sivatherium*, in the context of their anatomical traits, would help reveal how limb proportions and locomotor constraints may have evolved in the giraffid clade, and how similar constraints may have evolved in other tall animals, such as sauropod dinosaurs.

Here we question whether elongate, cursorial limbs constrain locomotion, rather than facilitate it. Our first prediction is that giraffes’ EMA is lower than expected for an animal of large body mass. To address this prediction, we used a synthesis of experimental data and musculoskeletal modelling to compare EMA of the giraffe forelimb (taken as the mean of EMA values at each joint) during walking to EMA values for animals ranging from mice to horses. Previous experimental work has demonstrated that forelimb and hindlimb EMAs in quadrupedal mammals are comparable (8, 11). We also use these data to test if low EMA may result in greater locomotor cost in giraffes by estimating active muscle volumes required during stance phase (9-11). Our second prediction is that EMA in the giraffid clade is associated with the lengths and proportions of the limb, i.e. taxa with longer limbs have poorer leverage against GRFs. EMA throughout the stance phase was estimated using skeletal models of *Giraffa, Okapia* and *Sivatherium* forelimbs, with modelled kinematics and GRFs. *Okapia* was assumed to be representative of giraffids’ ancestral condition.

## Results

### Giraffe EMA

EMA values for each forelimb joint in the giraffe are displayed in Figure 3A. Mean EMA_imp_ and EMA_40_ (± 1 standard deviation) values were 0.34 (± 0.05) and 0.29 (± 0.05) respectively, with no apparent relationship with speed. Although these were statistically different measurements (t test, p < 0.001), the difference in biological terms was negligible. EMA was typically low at the start and end of stance (Figure S1), although forces are also low during this time (3). EMA tended to abruptly rise to (and fall from) infinity during the stance phase, due to the GRF vector passing through some joints’ centers of rotation.

**Figure 3.**
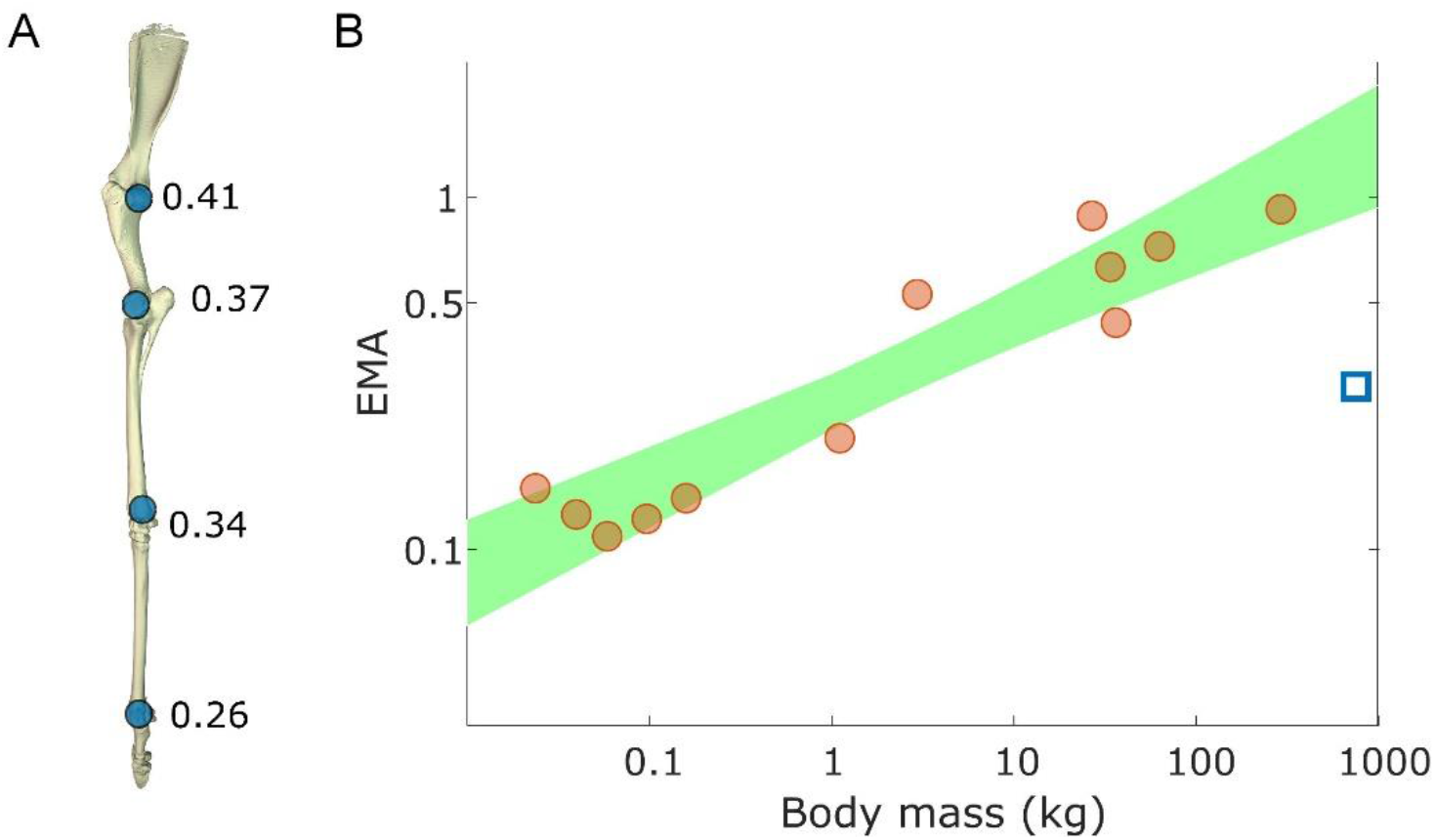
(A) Mean values of EMA for each joint of the giraffe forelimb (shoulder to MCP; shown in vertical reference pose). (B) Giraffe forelimb EMA (blue square) fell below the 95% prediction interval (shaded area), indicating that walking giraffes significantly deviate from the pattern seen in mammals of 0.03 – 297 kg at their trot-gallop transition (8).

EMA_40_ was compared with data from other mammalian quadrupeds. Using the comparative dataset of animals ranging from 0.024 to 297 kg (20), an animal with body mass 780 kg was predicted to have an EMA of 1.3 (with 95% prediction interval 0.88 – 1.93). Giraffe forelimb EMA falls well below the 95% prediction interval (Figure 3B); about 24% of predicted EMA.

EMA_imp_ sensitivity varied with the magnitude of COP displacement in *Giraffa* (Figure S2). Displacement of the COP from its initial location at the distal third phalanx resulted in modest variation in EMA. Changes of this magnitude (or other plausible COP assumptions) did not alter the result that giraffes’ EMA falls well below the scaling prediction for smaller mammals. Estimated active muscle volume for each trial ranged 40 – 89 cm^−3^ kg^−1^ m^−1^, with mean 54 (±14), and showed no apparent relationship with speed or stance duration.

### Comparisons of EMA between giraffids

We modelled the stance phase of *Giraffa, Sivatherium* and *Okapia* (Videos S1-3), using statically posed skeletal models, animated with experimental kinematics. We tested for any difference between this method (EMA_stat_) and the experimentally derived giraffe data (EMA_imp_). There was no statistical difference between the two methods (t test, p=0.26). We further checked for errors in modelled GRF moment arms and muscle moment arms, in case concurrent errors were effectively cancelling each other out, resulting in net agreement.

Mass and inertial properties were ignored in the static models, where EMA_stat_ was purely a geometric calculation (Eqn 1). This was a potential source of discrepancy when comparing with experimentally derived EMA_imp_, which did take these parameters into account (Eqn 4). To ensure we made sufficiently valid comparisons, we repeated EMA_imp_ measurements using a giraffe musculoskeletal model with all mass properties set to zero, which in effect was equivalent to the simple measurement of r/R. We found that EMA_imp_ for each trial was similar whether the limb’s mass properties were enabled or ignored (t test p = 0.065; Figure S3).

Other potential sources of error from the EMA_stat_ models for *Giraffa* included inaccurate muscle moment arm and/or GRF moment arm estimates. To test this, GRF moment arms were compared from the static models with moment arms from the inverse dynamics method (Figure S4). The GRF moment arms from the two methods, summarized as the mean moment arm, had a root mean square error (RMSE) of 6%. Therefore, we consider variable muscle moment arms to be the source of disparity between experimentally derived and modelled EMA.

The muscle moment arms measured from the static giraffe model were compared with the weighted mean moment arms (derived from the musculoskeletal model) used to calculate EMA_imp_ (Figure S5). The largest disparities were observed at the shoulder joint, where the extensor moment arm was over-estimated by 0.04 m (~67%); a result which led to a greater EMA value and a non-significant bias against our assumption that static and dynamically modelled moment arms were similar. We assumed that similar disparities in all three taxa likewise were non-significant, but not problematic for addressing our study’s key questions.

*Giraffa* incurred the greatest absolute GRF moment arms, followed by *Sivatherium* and *Okapia*, respectively (Figure 4A, S6). The muscle moment arms, modelled as the parasagittal distance from the estimated joint center of rotation to the bone surface, and normalized by shoulder height, were also compared. In most cases, *Sivatherium* had the largest muscle moment arms, with the exception of the MCP flexor moment arm (Figure 4B). There was imprecision associated with the measurement of the MCP moment arm in *Sivatherium*, as the proximal sesamoid bones were modelled and scaled from *Giraffa* (19). In most cases *Giraffa* possessed the smallest muscle moment arms. The greatest difference in muscle moment arms was between the *Giraffa* and *Sivatherium* olecranon process at the elbow joint.

**Figure 4.**
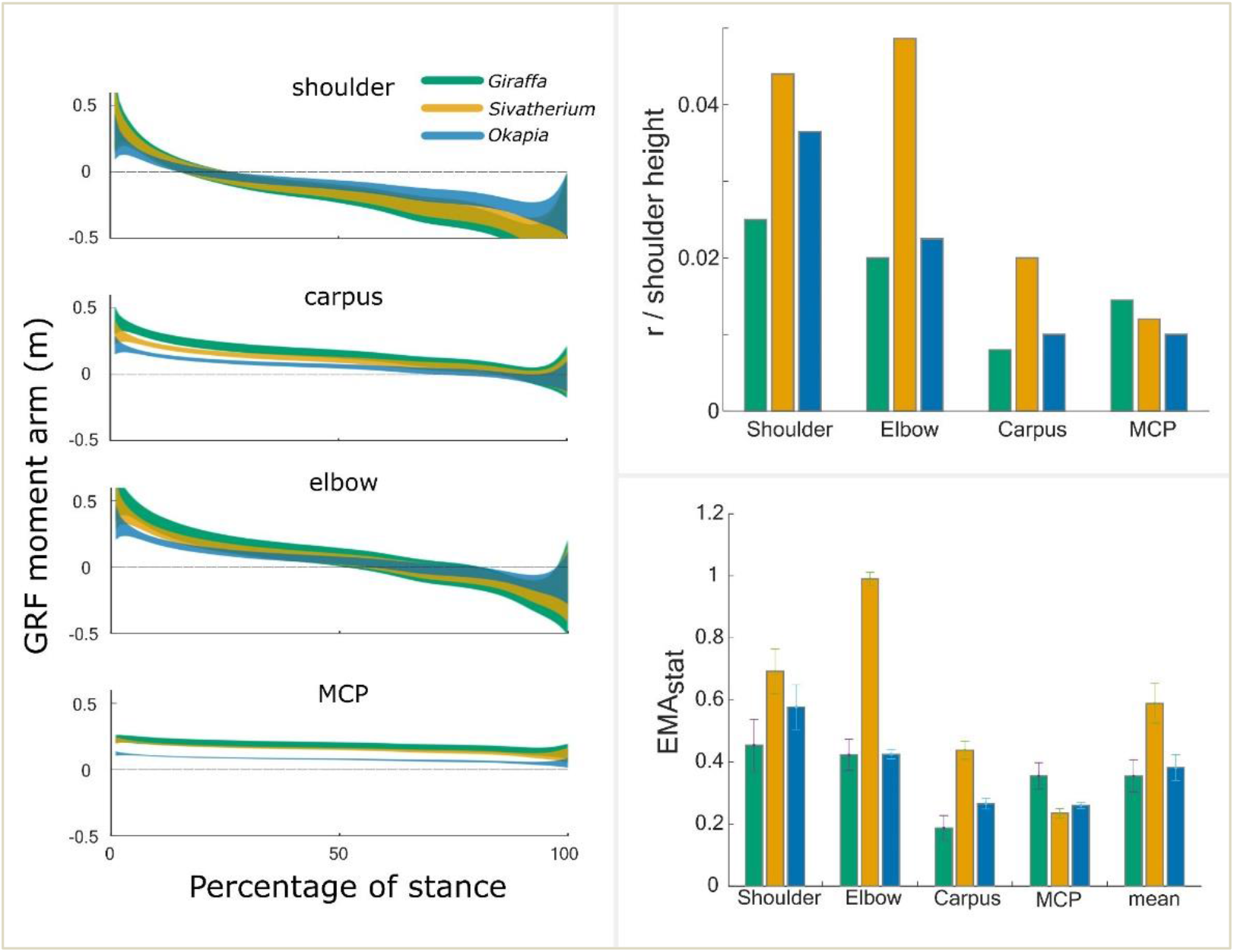
(A) Modelled GRF moment arms in three giraffids, derived using data from 14 experimental trials from *Giraffa*. Shaded regions show 95% confidence intervals for mean moment arm at each timepoint. *Giraffa* consistently had the greatest magnitude GRF moment arms. (B) Estimations of normalized muscle moment arms for the shoulder extensors, elbow extensors, carpal flexors and MCP flexors (i.e. antigravity muscles used to calculate EMA). (C) EMA_stat_ throughout the stance phase. Due to a combination of large GRF moment arms and modest muscle moment arms, *Giraffa* incurred the lowest EMA of the giraffids studied. Error bars denote 1 standard deviation.

The GRF and muscle moment arms above were used to estimate EMA_stat_ over the course of a modelled stance phase for the three giraffid models. *Sivatherium* was estimated to have the greatest EMA_stat_, followed by *Okapia* and *Giraffa* (Figure 4C).

## Discussion

### EMA in giraffes

Giraffes have a smaller than expected EMA for an animal of such large body mass (Figure 3B). We found that a giraffe using a typical lateral sequence walking gait had a forelimb EMA_40_ of 0.29 rather than the value of 1.3 predicted from scaling of forelimb EMA in smaller taxa (8, 12, 20). We predict that the same conclusions can be applied to the hindlimb, which (as in other cursorial mammals) display similar patterns of EMA (8). This value is also less than half that for walking Asian elephants (*Elephas maximus*) (11), and unlike Asian elephants, the giraffe’s EMA did not change within the (narrow) range of observed speed.

We found that two common methods for calculating EMA (EMA_imp_ and EMA_40_) yielded similar results (t-test, p=0.26) and led to comparable conclusions. EMA_40_ in giraffes was outside of the 95% prediction interval of the log-transformed linear model from Biewener (2005) (Figure 3B), and was consistent with the concept that an EMA plateau exists in animals with body mass in excess of 300 kg (11, 13, 20). Reasons for low EMA values in *Giraffa* can be ascribed to the magnitudes of the GRF and/or muscle moment arms. With regard to GRF moment arms, animals larger than horses probably are unable to align their GRF vector even closer to their joint centers to minimize R and maximize EMA (21), via increased straightening of the limb. In the case of the giraffe, our comparisons between closely related giraffid species suggest that their long segment lengths and shoulder height (and thus “cursorial” limb morphology) predispose them to exaggerated GRF moment arms (Figure 4A).

Alternatively, animals may be able to counter large GRF moment arms with similarly large muscle moment arms. This does not appear to be the case for giraffes. For example, the shoulder extensor moment arm of the long head of the triceps brachii muscle was 0.10 m throughout stance, similar to the 0.13 m predicted for a 780 kg animal (22). The moment arms of giraffes’ major muscle groups are summarized in Table S1. We surmise that giraffes are ill-equipped to effectively offset such large GRF moment arms, resulting in low EMA.

Since the calculation of EMA dictates that it is inversely proportional to the active muscle volume (11), giraffes’ relatively small EMA during walking suggests that a large volume of muscle is recruited to oppose the GRFs that act on a limb. Surprisingly though, giraffes’ mass-specific muscle volume recruitment (V_musc_; 40 – 89 cm^−3^ kg^−1^ m^−1^) during walking is 4 – 8 times larger than in walking humans, but broadly in line with other quadrupeds, including dogs, quadrupedal chimpanzees and elephants (11, 23). Low EMA is instead compensated for by long step lengths and relatively short muscle fascicles; shorter than the predictions for other non-hopping mammals (22).

We were unable to correlate active muscle volume with metabolic cost of transport in walking giraffes, as such data are unavailable. But since active muscle volume is correlated with metabolic costs in birds and mammalian quadrupeds and bipeds (11), we similarly expect that giraffes incur modest cost of transport at the slow walking speeds observed, and speculate that locomotor economy is an important factor in determining preferred speed. We previously suggested that giraffes avoid speeds outside of this optimum, due to sharp increases in metabolic cost (3). We predict that faster speeds during walking or their galloping gait (4) are met with increased step lengths and (potentially) changes in limb EMA, leading to higher metabolic costs, and that this places a constraint on giraffes’ athletic performance.

EMA also relates to mechanical stress of supportive tissues. The scaling of EMA α BM^0.26^ in mammals from 0.03 – 300 kg BM, combined with PCSA α BM^0.80^, suggests that supportive tissue stresses are nearly independent of body mass (20, 22). As a consequence, animals with below-expected EMA may risk higher skeletal and muscle stress, and catastrophic failure if no other changes are made to their locomotor dynamics. In order to reduce the risk of tissue failure, giraffes should be forced to reduce their athletic ability (7). Low EMA may explain giraffes’ limited capacity for speed (4, 24, 25), and may be a contributing factor as to why giraffes do not gallop in a dynamically similar manner to other mammalian quadrupeds (4).

We reject the notion that giraffes’ extreme height disposes them to a ‘mechanical advantage’ in locomotion (1), or that their long limbs facilitate fast speed locomotion (2). Instead, we find support for our prediction that extreme height and limb length in animals such as giraffids exceeding 300 kg results in increased GRF moment arms, and logically, reduced EMA.

### EMA of giraffid species

EMA_stat_ from *Giraffa, Sivatherium* and *Okapia* - three phenotypically distinct giraffids - were estimated, using statically posed skeletal models. We used this modelling method to predict how changes in limb segment lengths can alter EMA_stat_ of a limb, and as a consequence, drive changes in locomotor behaviour. At each joint, *Giraffa* consistently had the greatest absolute GRF moment arms (and lowest EMA_stat_), contrasting with *Okapia* which had the smallest (Figure 4A, S8). When these moment arms were normalized to shoulder height, these differences disappeared. This is consistent with the assumption of geometrically similar GRF orientation between the three studied taxa, and implies that GRF moment arms should scale isometrically with shoulder height. If this assumption is experimentally confirmed for a phylogenetically diverse sample of cursorial mammals, tall animals will be subject to large GRF moment arms (Figure 5); this offers an explanation as to why EMA diminishes in mammals exceeding horse size.

**Figure 5.**
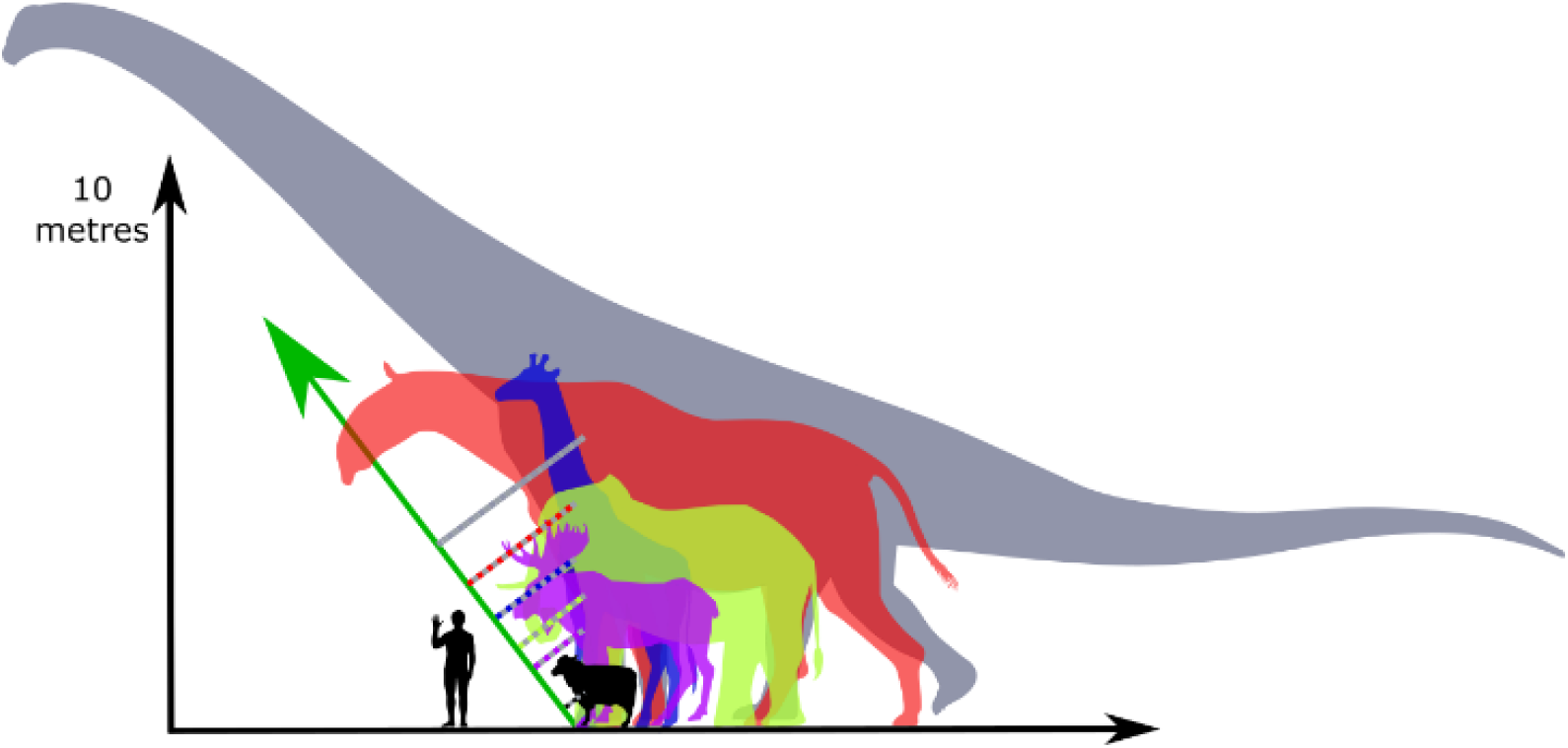
At increasing limb length, and given consistent GRF orientation (green arrow) and limb posture, GRF moment arms (dotted lines) are predicted to increase, resulting in progressively reduced EMA. In ascending order of size: *Ovis aries*, *Alces alces*, *Elephas maximus*, *Giraffa camelopardalis*, *Paraceratherium transouralicum*, *Patagotitan mayorum*. Image adapted with permission from work by Wikipedia artist Steveoc 86 and https://www.freepik.com/macrovector.

EMA is also dependent on muscle moment arm length. To test whether or not large (>300 kg) body mass is strictly associated with low EMA_stat_, we modelled the muscle moment arms and GRF moment arms of *Sivatherium giganteum*. Despite sharing a similar body mass, and probably a similarly upright limb posture (Figure 1B), mean EMA_stat_ was predicted to be 2 times greater in *Sivatherium*, compared with G*iraffa*. The source of this apparent difference lay both in the differences in GRF moment arm (Figures 4A, 5) and *Sivatherium’s* relatively large ‘antigravity’ muscle moment arms (Figure 4B).

The robustness of the *Sivatherium* skeleton is exemplified by the olecranon process of the fused radioulna bone, which is a useful proxy for the magnitude of the elbow extensor muscles’ moment arm. The ‘considerable’ projection of the olecranon was noted in an early fossil description (26). The olecranon process of *Sivatherium* was indeed considerably longer than in *Giraffa* (Table S1), by 0.07 m (an 80% difference in parasagittal length, despite similar body mass). Hence we speculate that *Sivatherium* was better equipped to offset the GRF moment arms encountered during the stance phase, than the more gracile *Giraffa*.

We surmise that giraffes’ extreme height has incurred a locomotor performance penalty, which may reflect their relatively modest athleticism (25). This complements the specializations in behavior and ecology seen in megaherbivores (27). For example, reduced predation in adult giraffes (28, 29) may relax the selection pressures for high performance traits, such as speed and endurance. Such relaxation of selection pressures may subsequently facilitate the expression of novel or extreme morphology.

## Conclusions

We have highlighted that giraffes use lower than expected effective mechanical advantage, as their musculoskeletal morphology (such as the ulna’s olecranon process) is insufficient to maintain the observed trend in EMA in animals up to 300 kg. Our results from an analysis of modelled GRF moment arms and muscle moment arms suggested that giraffes’ EMA is similar to okapis, a giraffid with lower body mass and more plesiomorphic locomotor traits. Low EMA was not ubiquitous among the giraffids, as *Sivatherium giganteum* was predicted to have greater EMA; but still low compared to smaller mammals, even horses. The differential EMA between *Sivatherium* and *Giraffa* may reflect behavioural or athletic differences between these two similarly sized giraffids. Whilst giraffes’ feeding ability is driven by extreme height, it appears that this specialization has come with a functional trade-off with locomotor performance.

## Materials and Methods

### Dynamic musculoskeletal modelling

A rigid-body giraffe musculoskeletal model was developed using the software package Software for Interactive Musculoskeletal Modeling (SIMM v6.0; MusculoGraphics Inc, California, USA), as follows. The skeleton of a cadaveric forelimb from a captive bred 7 year old male giraffe donated postmortem by a local zoo, with body mass 880 kg, was segmented from CT images (2.5 mm slice thickness, 100 kV, 200 Ma, Lightspeed Pro 16 slice CT, GE Medical, Buckinghamshire, UK), and the resulting meshes exported as .stl files using the software package Mimics (v19.0 Materialise, Leuven, Belgium). The digitized bones of the forelimb were then used to construct a model (Figure 2B) consisting of five body segments (scapula, humerus, radioulna, metacarpus and phalanges). Joint axes were assigned, and the limb segments were aligned into a neutral reference pose (all joints at 0° = vertically aligned) using the software Maya (2016, Autodesk, California, USA). Joint axes were restricted to flexion and extension (i.e. hinge joints).

Muscle paths were added in SIMM, following established methods (30-32), guided using muscle geometry derived from CT data and gross dissection of the cadaver. The origins of forelimb extrinsic muscles were estimated in the model, as cadaveric geometry for the neck and skull were unavailable. Thirty-one musculotendon actuators were included (Supplementary Information). The mass and centre of mass (COM) of each segment (including soft tissues) were estimated with the methodology of (33) and (34), where the convex hull and subsequent mass parameters for each segment were calculated using the convex hull function of Meshlab version 2016.12 (35) and custom code written in Matlab (Mathworks, Massachusetts, USA). The geometry of the 880 kg giraffe model was isometrically scaled to the size of a 780 kg giraffe using OpenSim 3.3 (36), to match data from an experimental subject.

The calculation of EMA in Eqn. 1 is derived from the notion that joint moments induced by a GRF must be balanced by an opposing and equal muscle moment, i.e.:

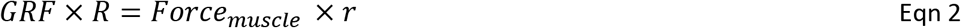

Rearranged, EMA can be expressed both in terms of moment arms and in terms of forces:

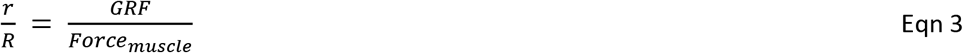

Forces can be considered over the duration of the stance phase by calculating impulses (force-time integrals). In this way, EMA can be expressed as:

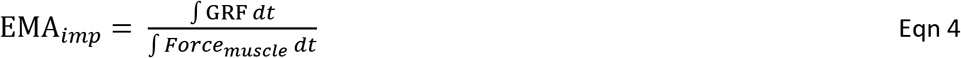

Using the impulses (9, 11, 37) has the advantage that the entire stance duration can be considered, not just a single instant or the mean across a step. Overall limb EMA_imp_ was calculated as the mean of EMA_imp_ at each joint (12).

Experimentally derived GRF and kinematic data (3) were used to calculate EMA at each joint, throughout the stance phase. Briefly, three adult reticulated giraffes walked over a three-axis force platform, in front of a video camera (Video S1). Joint centers were visually estimated and digitized using DLTV6 (38). 14 walking steps from one individual were selected from the larger dataset, with speed ranging from 0.8 to 1.2 ms^−1^ (0.04 to 0.08 Froude number). These were selected on the basis that the giraffe was not obscured by any foreground objects. This work was conducted with ethical approval (number URN 2016 1538) from the Clinical Research Ethical Review Board of the Royal Veterinary College, University of London.

Forces (e.g. of muscles acting around a joint) can be estimated from moment and muscle moment arm (Eqn. 3), assuming static equilibrium:

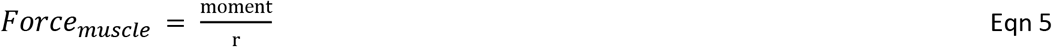

Total net moments acting at each joint were calculated using the inverse dynamics function in OpenSim 3.3 (36), where inertial (M_inert_) and gravitational (M_grav_) moments at the shoulder, elbow, carpus and MCP were considered along with the moments required to generate ground reaction force (M_GRF_)(9). The integral of total muscle force acting around each joint (i.e., *Force_muscle_* in equation 4) was calculated by dividing joint moments by the weighted mean muscle moment arm for muscles crossing that joint (equation 5 and see below). When a joint had variable action during stance (e.g. flexion followed by extension), force integrals for flexion and extension were separately calculated using their respective moment arm, and then summed to give total force.

The agonist muscle moment arm (r, Figure 2A) for each joint was calculated as the mean moment arm of the muscles at the time of peak GRF, weighted by each muscle’s contribution to total muscle physiological cross-sectional area (PCSA; see below), and with the numerical subscripts for r and PCSA below referring to each muscle’s moment arm or PCSA. This assumed that all agonist muscles were similarly active (9, 12) (Eqn 6). We did not address the issue of co-contraction by antagonist muscle groups, as these forces were assumed to be non-significant with respect to total muscle force. This approach keeps our analysis maximally comparable to other studies of mammalian EMA, vs. a more comprehensive dynamic simulation analysis.

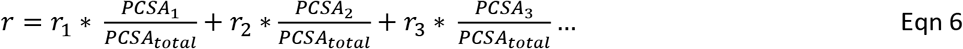

PCSAs of muscles from the same 880 kg individual were measured using muscle architecture methods from muscle mass, pennation, and mean fascicle length (37). The extrinsic muscles of the adult forelimb were missing; the PCSAs of these muscles were estimated by isometrically scaling PCSA of the corresponding muscles from a sub-adult giraffe cadaver, with body mass 480 kg. Isometry was chosen as an assumption in the absence of other data, as the bones of the forelimb scale with or close to isometry in the post-natal giraffe (39). Modest allometry of these missing muscles would not be expected to influence our results or conclusions in a pronounced way.

Whilst recent studies have used the above impulse method to calculate EMA (9, 11, 40), EMA from a varied range of mammalian species has been previously calculated as the mean ratio r/R, during the middle third of stance and at the trot-gallop transition (12). To facilitate comparisons between giraffes and other terrestrial mammals, EMA was additionally calculated in a more comparable manner. For each joint, following Biewener (1989), r/R was calculated when M_GRF_ > 40% of maximum M_GRF_, which approximately corresponds to the middle third of stance. A mean value of EMA at each joint was calculated from this sample, here referred to as EMA_40_.

Giraffe forelimb EMA_40_ was compared with a compiled dataset of EMA from 12 other mammalian species (8). Data points from a logarithmic scatter plot from this publication were digitised and replotted. The data were log-transformed, and a least squares regression model was used to calculate the 95% prediction interval for the EMA versus body mass relationship. Following prior studies and considering the modest sample size, potential biases incurred by phylogeny were not addressed. All data were analysed using Matlab.

EMA calculations are sensitive to the location of the center of pressure (COP). COP data derived from raw force plate outputs in giraffes were excluded from this analysis due to excessive signal noise. In our model, the COP was fixed at the distal tip of the third phalanx. Placing the COP at this location facilitates repeatability of the method with different model taxa, but experimental data from a variety of animals show that COP is dynamic during the stance phase; tending to track cranially from an initial caudal position at the heel (41-44). A sensitivity analysis was performed to assess the effect of COP location on EMA for one trial, where the COP was randomly displaced (using Matlab) from the distal tip of the foot 100 times, to a maximum of 0.1 m (i.e. the length of the distal phalanges). EMA was then calculated in each case.

We estimated the mass-specific volume muscle activated per distance travelled for each of the trials (11, 23, 37), calculated as:

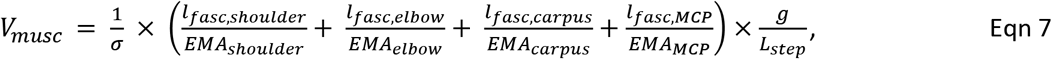

where V_musc_ is in units cm^−3^ kg^−1^ m^−1^, σ is assumed constant muscle stress (20 Ncm^−2^), *g* is acceleration due to gravity (9.81 ms^−2^), *l_fasc_* values are the mean agonist muscle fascicle lengths (in cm) at each joint, weighted by each muscle’s relative PCSA (similar to equation 6), EMA is derived from the ratio of GRF to muscle force (equation 4), and L_step_ is the horizontal distance travelled by the center of mass during the stance phase.

### Static musculoskeletal *modelling*

We generated biomechanical models of the forelimb stance phase for the extinct *Sivatherium giganteum* and the extant *Okapia johnstoni* to estimate EMA in these taxa. We chose the simplified approach of modelling the limbs as rigid multi-segmented structures. These models are termed ‘static’ because the internal joint angles were fixed; not driven by experimental kinematic data as for *Giraffa*; although all three taxa studied were analyzed using lever mechanics. The static models were used to estimate the GRF and muscle moment arms throughout stance (Figure 1B), during a modelled walking step. The model for *Okapia* was derived from photogrammetry of a complete mounted skeleton (specimen USNM 399337, Smithsonian Institution, Washington, DC, USA), mounted in a standing posture. A 3D mesh was generated from 300 digital photographs of the specimen using Photoscan v1.4 (Agisoft, St.Petersburg, Russia) and Meshlab v2016 (35). The forelimb skeleton of *Sivatherium giganteum* was reconstructed from ten fossil specimens from the Natural History Museum, UK (Table S2). 3D surface meshes were derived from photogrammetry of these specimens, and articulated into a reconstruction. It is likely that these post-cranial specimens may be attributed to the same individual (45). The missing distal phalanx and proximal sesamoid bones were scaled from the same 880 kg giraffe (19).

Stance phase postures and all measurements were implemented in Maya. Mid-stance forelimb joint angles for the okapi (Table S3) were derived from walking in healthy okapis (personal communication [46]). A reconstruction of the *Sivatherium* mid-stance posture required three joint angles to be assumed, for the elbow, carpus and metacarpophalangeal (MCP) joints. The elbow angle was estimated by positioning the olecranon process of the radioulna perpendicular to the long axis of the humerus (47). The carpal joint angle was assumed to be fixed in a neutral position (0°) during stance. There is no tested method to predict MCP joint angle in extinct species using surface bone geometry. Thus for the current purpose, we speculated that loading at the MCP joint, due to body weight, was similar in *Sivatherium* as it is in *Giraffa*, given their similar body masses (19). We therefore assigned the same internal MCP angle to *Sivatherium*, as for the mid-stance giraffe model.

To model limb joint kinematics during stance, each limb was modelled as a stiff inverted pendulum (48), whereby the rigid limb vaults over a pivot. The most distal extremity of the third phalanx was assumed to be the rotation point. The angular sweep of the forelimb about this point was modelled on the motion of the giraffe’s shoulder through a walking stance phase. The unit vector of the shoulder position (from the toe) was measured at each timestep throughout stance, and imposed on the models of *Sivatherium* and *Okapi*. It was reasonable to extrapolate *Giraffa* kinematics to closely related species, considering that giraffes walk in a dynamically similar fashion to other mammalian quadrupeds (3), and more specifically similar to other cetartiodactyls ranging in size from domestic sheep (*Ovis aries*) to giraffes (49).

Model GRF vectors were required for the extinct giraffid *Sivatherium*, and for *Okapia*. *Giraffa*, as a closely related species, was used to model the GRFs of *Sivatherium* and *Okapia*. The validity of this approach was tested by comparing the GRF unit vectors of giraffes with other cetartiodactyl ungulates. During a steady state walking step, the unit GRF vector changes from positive (deceleration) to negative (acceleration). To assess whether the GRF vector is consistent amongst different mammalian cursorial taxa, the unit vectors of a giraffe were compared with two other ungulates whose phylogenetic relationships form a close bracket around the position of *Giraffa* (50). If a trait is conserved within this bracket (in this case a postural trait, supported by relatively conservative morphology), it can be assumed that all descendants of the root ancestor (including *Okapia* and *Sivatherium*) similarly share this character (51, 52). The unit vectors from the walking gait of red deer (*Cervus elephas*) and dromedary (*Camelus dromedarius*) were collected using the same force plate equipment (53), and compared with the stance phase unit vectors from the giraffe. Their GRF unit vectors showed a consistent pattern of change (Figure S7), and fall within the giraffe inter-trial variation.

GRF moment arms (R) with respect to the shoulder, elbow, carpus and MCP joint were calculated from the toe – joint vector (a) and GRF vector (b):

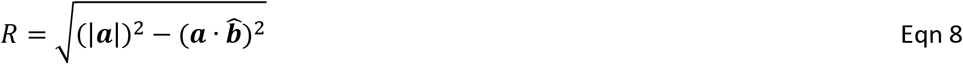

Muscle moment arms (r) were simplified to a single measurement of the flexor moment arm at the carpus and MCP joint, and extensor moment arms at the shoulder and elbow (Figure S8). EMA_stat_ was calculated as r/R (Equation 3) at each percentage time step during stance. Only flexor muscle moment arms at the carpus and MCP joints were included in the analysis, as these account for the anti-gravity function throughout the stance phase. In the case of the shoulder and elbow, the flexor muscle moment arms depend on prior interpretations of muscle origins and insertions (i.e. a musculoskeletal model), and were not included in this analysis (Figure S1). Adopting this approach permitted readily objective comparisons between specimens including the fossil giraffid. We then compared these simplified geometric measurements in the static *Giraffa* model with those derived from experimental inverse dynamics, to assess the validity of this approach.

This static modelling approach made the following assumptions that throughout the stance phase: (1) GRF unit vectors are the same in *Giraffa, Okapia* and *Sivatherium;* (2) the toe to shoulder unit vectors are the same in *Giraffa, Okapia* and *Sivatherium;* (3) joint angles are constant throughout stance. These assumptions are static simplifications of an otherwise dynamic behavior. In order to assess the validity of the subsequent EMA calculations, an additional *Giraffa* static model was created using the same methodology. The static model’s moment arms and EMA were compared with those derived from the experimental data (Figure S1, S4-5).

## Acknowledgements

We thank Pip Brewer from the Natural History Museum (UK) and Darrin Lunde from the Smithsonian National Museum of Natural History (USA) for access to giraffid specimens; Vivian Allen, Jeffery Rankin and Heinrich Mallison for helpful discussions and technical technical assistance; and Kristiaan D’Août for data on okapi kinematics. Ashleigh Wiseman and Peter Bishop provided helpful comments on the manuscript. Thank you also to Peter Aerts and Helen Birch for their feedback on CB’s PhD thesis chapter from which this publication is derived. We thank NERC for funding this work (PhD studentship for C.B.; grant no. NE/K004751/1 to J.R.H.).

## References

1. G. Mitchell, J. Skinner, On the origin, evolution and phylogeny of giraffes Giraffa camelopardalis. Transactions of the Royal Society of South Africa 58, 51–73 (2003).

2. C. Pincher, Evolution of the giraffe. Nature 164, 29–29 (1949).

3. C. Basu, A.M. Wilson, J. R. Hutchinson, The locomotor kinematics and ground reaction forces of walking giraffes. Journal of Experimental Biology 222(2019).

4. C. K. Basu, F. Deacon, J. R. Hutchinson, A.M. Wilson, The running kinematics of free-roaming giraffes, measured using a low cost unmanned aerial vehicle (UAV). PeerJ 7, e6312 (2019).

5. T. Garland, Scaling the Ecological Cost of Transport to Body Mass in Terrestrial Mammals. The American Naturalist 121, 571–587 (1983).

6. M. R. Hirt, W. Jetz, B. C. Rall, U. Brose, A general scaling law reveals why the largest animals are not the fastest. Nature Ecology & Evolution 1, 1116 (2017).

7. T. J. M. Dick, C. J. Clemente, Where Have All the Giants Gone? How Animals Deal with the Problem of Size. PLOS Biology 15(2017).

8. A. A. Biewener, Biomechanical consequences of scaling. Journal of Experimental Biology 208, 1665–1676 (2005).

9. A. A. Biewener, C.T. Farley, T. J. Roberts, M. Temaner, Muscle mechanical advantage of human walking and running: implications for energy cost. Journal of Applied Physiology 97, 2266–2274 (2004).

10. H. Pontzer, Effective limb length and the scaling of locomotor cost in terrestrial animals. Journal of Experimental Biology 210, 1752–1761 (2007).

11. L. Ren, C. E. Miller, R. Lair, J. R. Hutchinson, Integration of biomechanical compliance, leverage, and power in elephant limbs. Proceedings of the National Academy of Sciences 107, 7078–7082 (2010).

12. A. Biewener, Scaling body support in mammals: limb posture and muscle mechanics. Science 245, 45–48 (1989).

13. J. R. Hutchinson, M. Garcia, Tyrannosaurus was not a fast runner. Nature 415, 1018–1021 (2002).

14. N. Solounias, Family Giraffidae. The evolution of artiodactyls, 257 (2007).

15. W. R. Hamilton, Fossil giraffes from the Miocene of Africa and a revision of the phylogeny of the Giraffoidea. Philosophical transactions of the Royal Society of London. Series B, Biological sciences, 165–229 (1978).

16. J. E. Fa, A. Purvis, Body size, diet and population density in Afrotropical forest mammals: a comparison with neotropical species. Journal of Animal Ecology, 98–112 (1997).

17. M. Danowitz, N. Solounias, The Cervical Osteology of Okapia johnstoni and Giraffa camelopardalis. PloS one 10, e0136552 (2015).

18. N. A. Badlangana, JW; Manger, P, The giraffe cervical vertebral column: a heuristic example in understanding evolutionary processes? Zoological Journal of the Linnean Society 155, 736–757 (2009).

19. C. Basu, P.L. Falkingham, J.R. Hutchinson, The extinct, giant giraffid Sivatherium giganteum: skeletal reconstruction and body mass estimation. Biology Letters 12(2016).

20. A. A. Biewener, Biomechanics of mammalian terrestrial locomotion. Science 250, 1097–1103 (1990).

21. J. R. Hutchinson, Response: Of ideas, dichotomies, methods, and data – how much do elephant kinematics differ from those of other large animals? Journal of Experimental Biology 212, 153–154 (2009).

22. R. Alexander, A. Jayes, G. Maloiy, E. Wathuta, Allometry of the leg muscles of mammals. Journal of Zoology 194, 539–552 (1981).

23. H. Pontzer, D. A. Raichlen, M. D. Sockol, The metabolic cost of walking in humans, chimpanzees, and early hominins. Journal of Human Evolution 56, 43–54 (2009).

24. T. Garland, C.M. Janis, Does metatarsal/femur ratio predict maximal running speed in cursorial mammals? Journal of Zoology 229, 133–151 (1993).

25. T. Garland, The relation between maximal running speed and body mass in terrestrial mammals. Journal of Zoology 199, 157–170 (1983).

26. H. Falconer, Palaeontological Memoirs and Notes of the late Hugh Falconer: With a Biographical Sketch of the Author Compiled and edited by Charles Murchison (Rob. Hardwicke, 1868).

27. J. E. Cohen, S. L. Pimm, P. Yodzis, J. Saldaña, Body sizes of animal predators and animal prey in food webs. Journal of animal ecology, 67–78 (1993).

28. R. Pellew, The giraffe and its food resource in the Serengeti. I. Composition, biomass and production of available browse. African Journal of Ecology 21, 241–267 (1983).

29. N. Owen-Smith, M. G. L. Mills, Predator–prey size relationships in an African large-mammal food web. Journal of Animal Ecology 77, 173–183 (2008).

30. J. R. Hutchinson, F. C. Anderson, S. S. Blemker, S. L. Delp, Analysis of hindlimb muscle moment arms in Tyrannosaurus rex using a three-dimensional musculoskeletal computer model: implications for stance, gait, and speed. Paleobiology 31, 676–701 (2005).

31. J. R. Hutchinson et al., Musculoskeletal modelling of an ostrich (Struthio camelus) pelvic limb: influence of limb orientation on muscular capacity during locomotion. PeerJ 3, e1001 (2015).

32. J. P. Charles, O. Cappellari, A. J. Spence, D. J. Wells, J. R. Hutchinson, Muscle moment arms and sensitivity analysis of a mouse hindlimb musculoskeletal model. Journal of Anatomy 10.1111/joa.12461 (2016).

33. W. Sellers et al., Minimum convex hull mass estimations of complete mounted skeletons. Biology Letters, rsbl20120263 (2012).

34. V. Allen, H. Paxton, J. R. Hutchinson, Variation in center of mass estimates for extant sauropsids and its importance for reconstructing inertial properties of extinct archosaurs. The Anatomical Record 292, 1442–1461 (2009).

35. P. Cignoni et al. (2008) Meshlab: an open-source mesh processing tool. in Eurographics Italian Chapter Conference, pp 129–136.

36. S. L. Delp et al., OpenSim: open-source software to create and analyze dynamic simulations of movement. Biomedical Engineering, IEEE Transactions on 54, 1940–1950 (2007).

37. T. J. Roberts, M. S. Chen, C.R. Taylor, Energetics of bipedal running. II. Limb design and running mechanics. Journal of Experimental Biology 201, 2753–2762 (1998).

38. T. L. Hedrick, Software techniques for two- and three-dimensional kinematic measurements of biological and biomimetic systems. Bioinspiration & biomimetics 3, 034001 (2008).

39. S. van Sittert, J. Skinner, G. Mitchell, Scaling of the appendicular skeleton of the giraffe (Giraffa camelopardalis). Journal of morphology 10.1002/jmor.20358 (2014).

40. A. D. Foster et al., Ontogeny of effective mechanical advantage in eastern cottontail rabbits (Sylvilagus floridanus). The Journal of experimental biology 222, jeb205237 (2019).

41. P. Van der Tol et al., The effect of preventive trimming on weight bearing and force balance on the claws of dairy cattle. Journal of Dairy Science 87, 1732–1738 (2004).

42. M. Grundy, P. Tosh, R. McLeish, L. Smidt, An investigation of the centres of pressure under the foot while walking. Bone & Joint Journal 57, 98–103 (1975).

43. O. Panagiotopoulou, T. C. Pataky, Z. Hill, J. R. Hutchinson, Statistical parametric mapping of the regional distribution and ontogenetic scaling of foot pressures during walking in Asian elephants (Elephas maximus). The Journal of experimental biology 215, 1584–1593 (2012).

44. D. R. Carrier, N. C. Heglund, K. D. Earls, Variable gearing during locomotion in the human musculoskeletal system. Science 265, 651–653 (1994).

45. H. Falconer, Palaeontological Memoirs and Notes of the late Hugh Falconer: With a Biographical Sketch of the Author Compiled and edited by Charles Murchison. II (Rob. Hardwicke, 1868).

46. K. D’Aout, C. Marien, K. Leus, P. Aerts (2005) Gait patterns and hoof impact in a captive giraffid, the Okapi (Okapia johnstoni). in COMPARATIVE BIOCHEMISTRY AND PHYSIOLOGY A-MOLECULAR & INTEGRATIVE PHYSIOLOGY (ELSEVIER SCIENCE INC 360 PARK AVE SOUTH, NEW YORK, NY 10010-1710 USA), pp S148–S148.

47. S.-i. Fujiwara, Olecranon orientation as an indicator of elbow joint angle in the stance phase, and estimation of forelimb posture in extinct quadruped animals. Journal of morphology 270, 1107–1121 (2009).

48. G. A. Cavagna, N.C. Heglund, C. R. Taylor, Mechanical work in terrestrial locomotion: two basic mechanisms for minimizing energy expenditure. American Journal of Physiology - Regulatory, Integrative and Comparative Physiology 233, R243–R261 (1977).

49. A. Pike, R. M. Alexander, The relationship between limb-segment proportions and joint kinematics for the hind limbs of quadrupedal mammals. Journal of Zoology 258, 427–433 (2002).

50. J. P. Zurano et al., Cetartiodactyla: Updating a time-calibrated molecular phylogeny. Molecular phylogenetics and evolution 133, 256–262 (2019).

51. L. M. Witmer, The extant phylogenetic bracket and the importance of reconstructing soft tissues in fossils. Functional morphology in vertebrate paleontology 1, 19–33 (1995).

52. H. N. Bryant, A.P. Russell, The role of phylogenetic analysis in the inference of unpreserved attributes of extinct taxa. Philosophical Transactions of the Royal Society of London. Series B: Biological Sciences 337, 405–418 (1992).

53. S. E. Warner et al., Size-related changes in foot impact mechanics in hoofed mammals. PloS one 8, e54784 (2013).

