## Supplemental figures and tables for "Low effective mechanical advantage of giraffes’ limbs during walking reveals trade-off between limb length and locomotor performance"

**This PDF file includes:**

Supplementary text

Figures S1 to S8

Tables S1 to S3

SI References

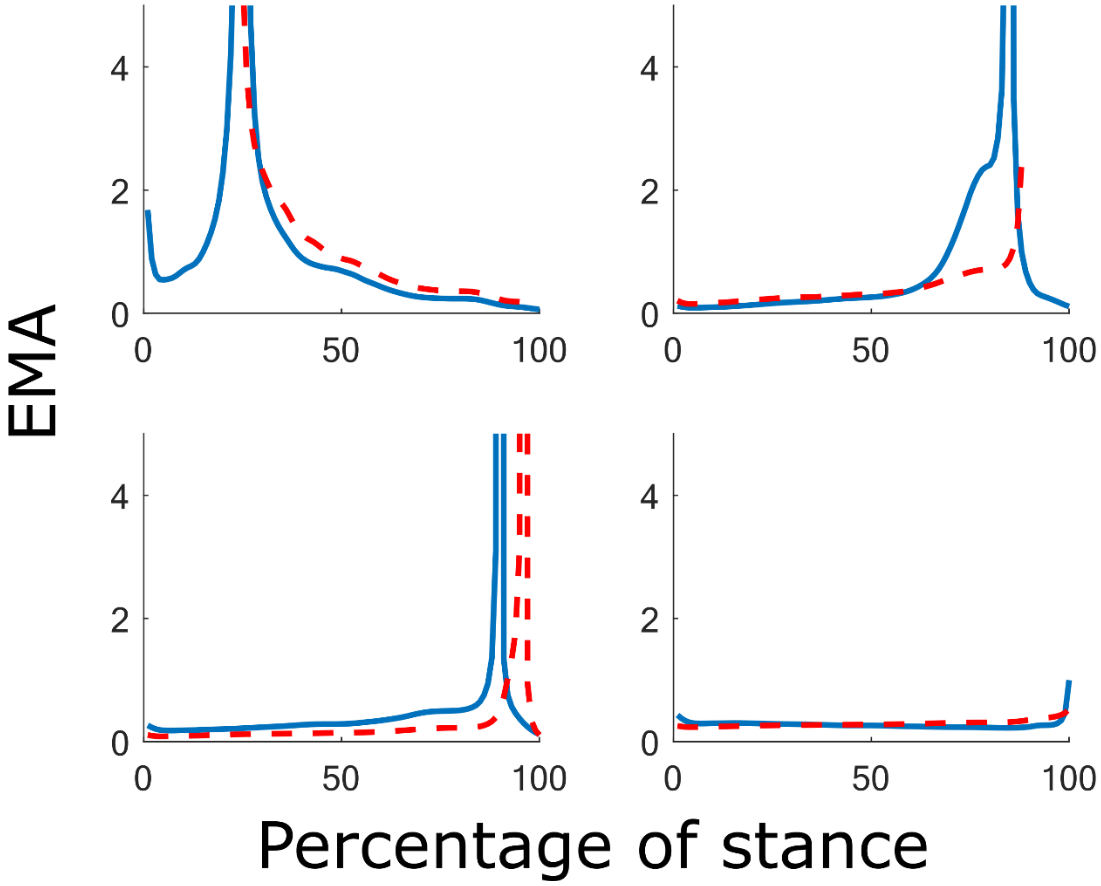


**Figure S1 Giraffe forelimb $EMA_{imp}$ derived from experimentally measured kinematics and kinetics (solid blue line) and inverse dynamics, from an exemplar stance phase, compared with $EMA_{stat}$; estimated from rigid skeletal models (red dotted line). Modelled $EMA_{stat}$ at the start of the shoulder timeseries and end of the elbow timeseries were not modelled, as the muscle moment arms during these times could not reliably be measured from skeletal specimens alone.**

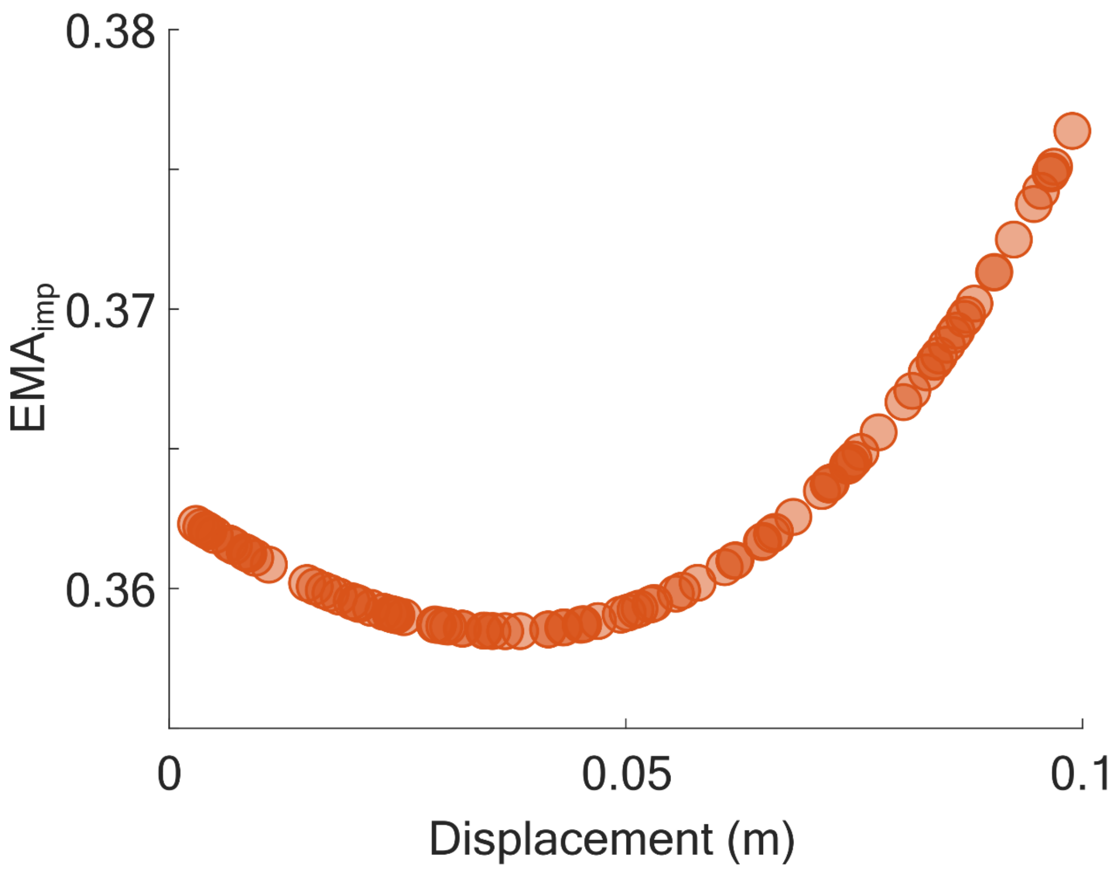


**Figure S2 EMA$_{imp}$ varied with centre of pressure location. Regardless of the location under the foot, EMA$_{imp}$ in the giraffe forelimb fell well below predictions for a large cursorial mammal (0.34 ± 0.05)**

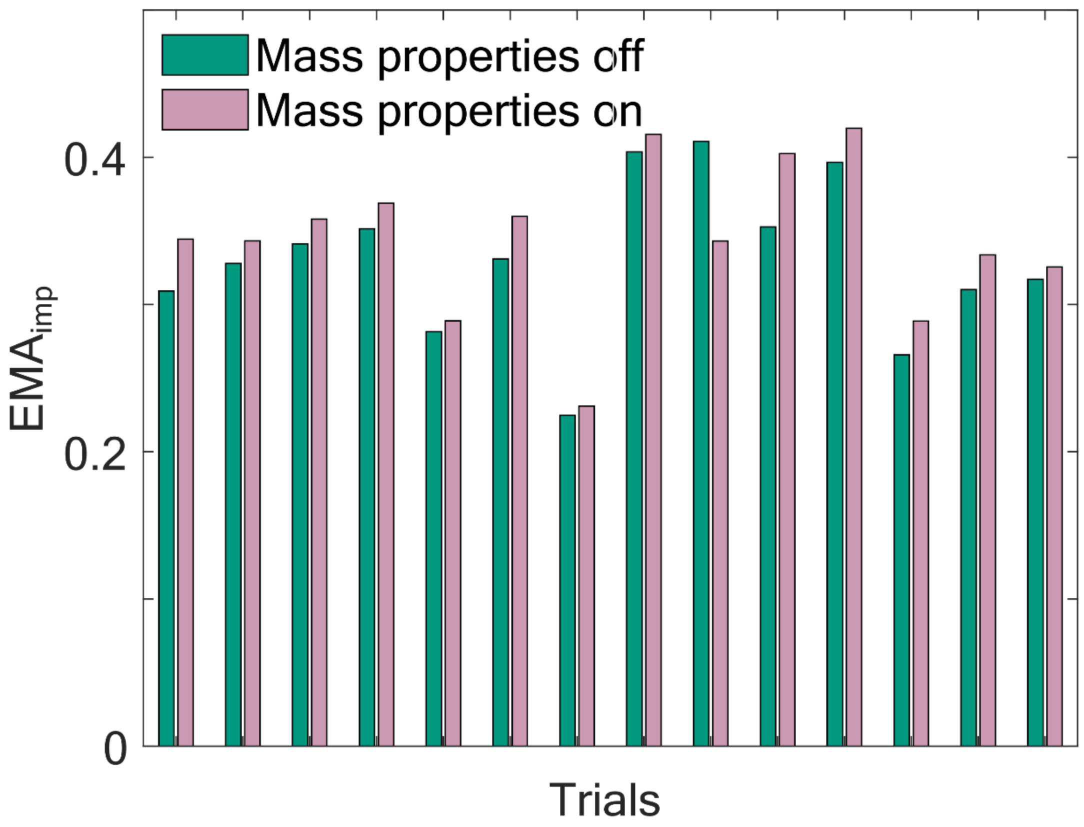


**Figure S3 Comparison of EMA$_{imp}$ derived using a massless musculoskeletal model using experimental data from 14 steps of an individual Giraffa, with a model featuring mass properties. EMA$_{imp}$ derived using inverse dynamics is comparable regardless of the model used (t-test, p = 0.26)**

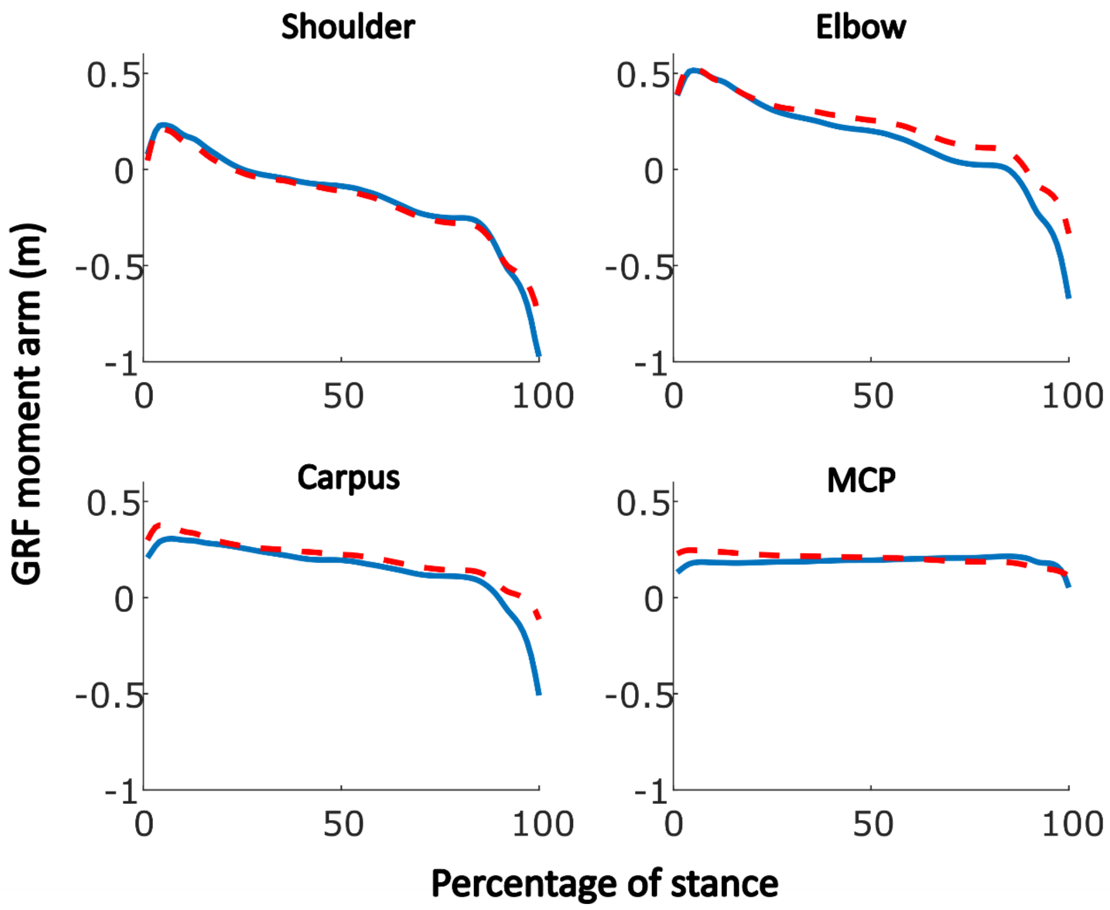


**Figure S4 The GRF moment arm (R), with respect to (A) shoulder, (B) elbow, (C) carpus and (D) the MCP joint in the giraffe. The solid blue line shows experimentally derived data from an exemplar trial, compared with the red dashed line which shows the moment arms derived from the static model. The two timeseries show comparable patterns of change, with minor error attributable to positional differences between the joint centres (experimental vs modelled). The GRF moment arms from the two methods, summarised as the mean, show a RMSE of 6%.**

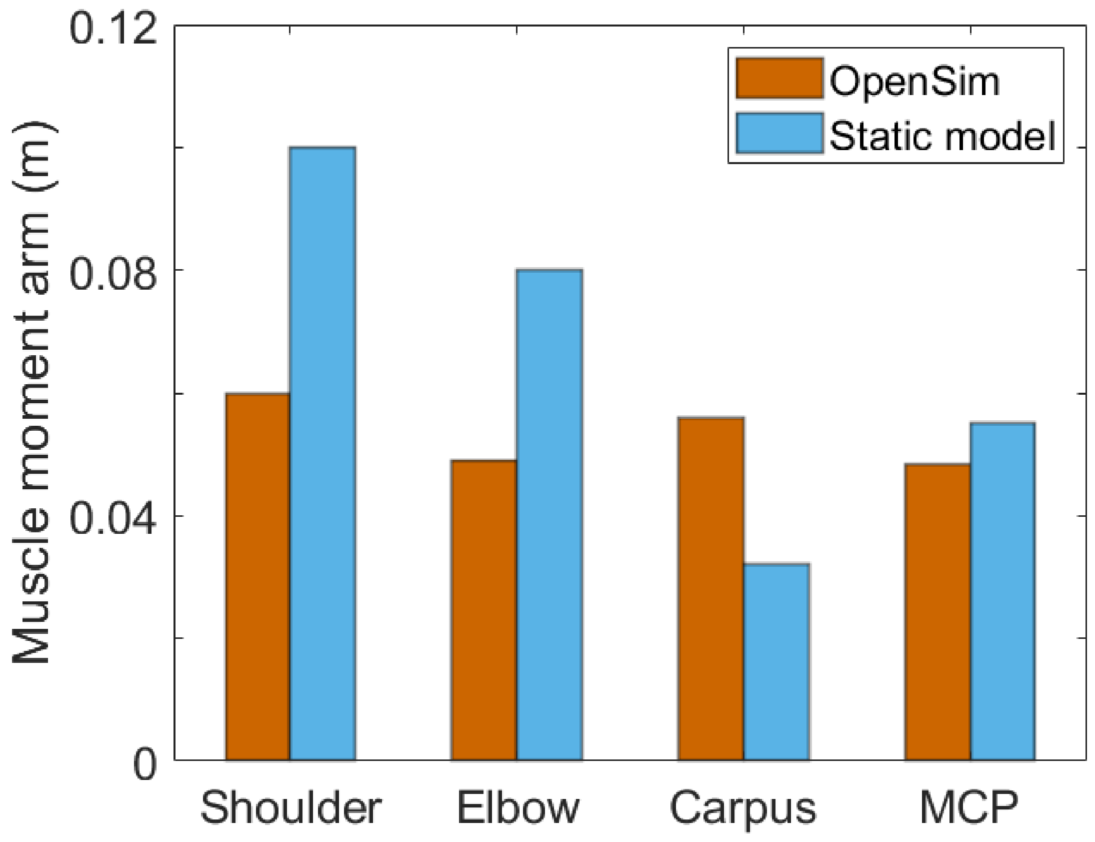


**Figure S5 Comparison of muscle moment arms (r) obtained from the giraffe static skeletal model (blue), compared with muscle moment arms exported from the OpenSim musculoskeletal model (orange). Bars refer to shoulder extensors, elbow extensors, carpal flexors, and MCP flexors. Overall, the static modelling approach tended to overestimate moment arms, especially for the shoulder.**

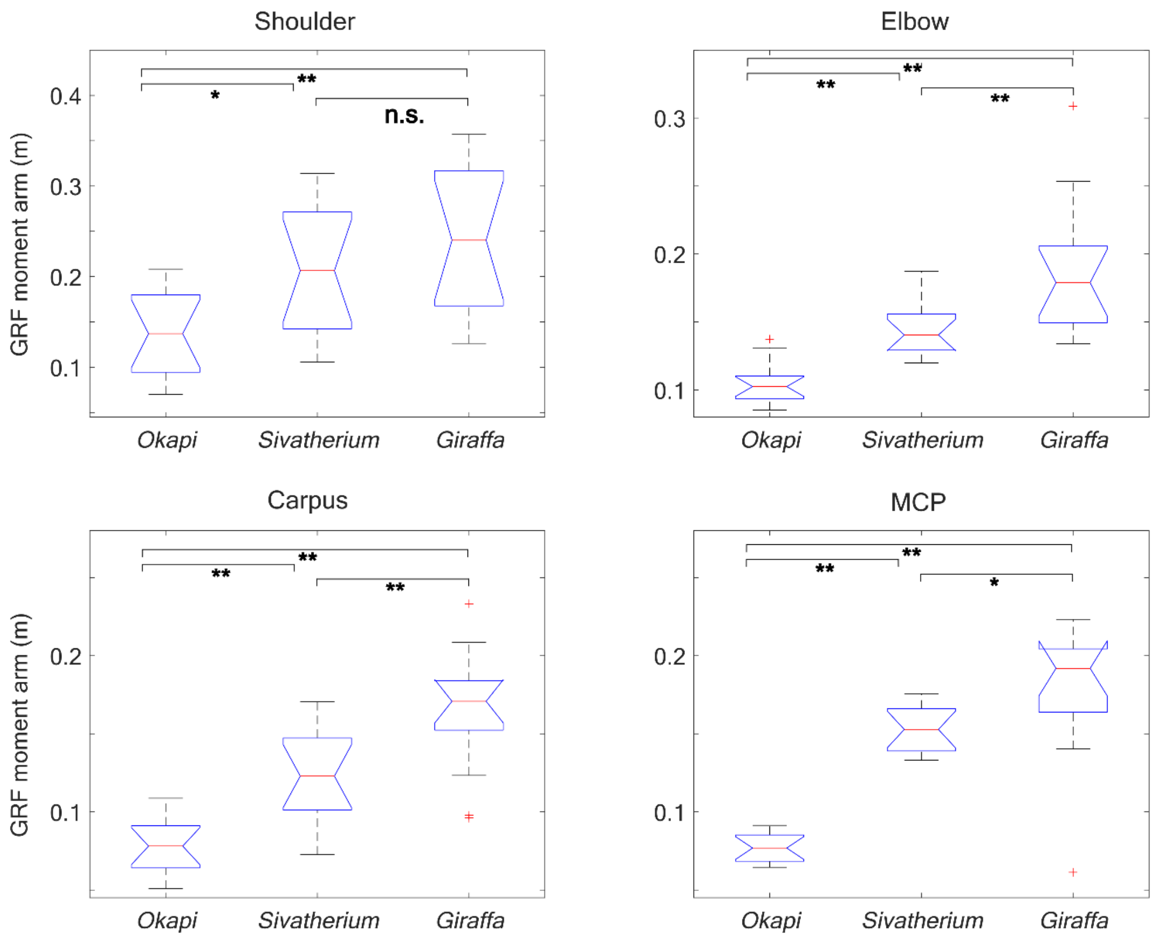


**Figure S6 Comparison of absolute GRF moment arms (R) between giraffids, expressed as the time averaged integral of the moment arms. Differences were evaluated by a one-way ANOVA, and are reported as not significant (n.s.), p<0.05 (*) or p<0.01 (**). These absolute GRF moment arms tended to be larger in larger giraffids.**

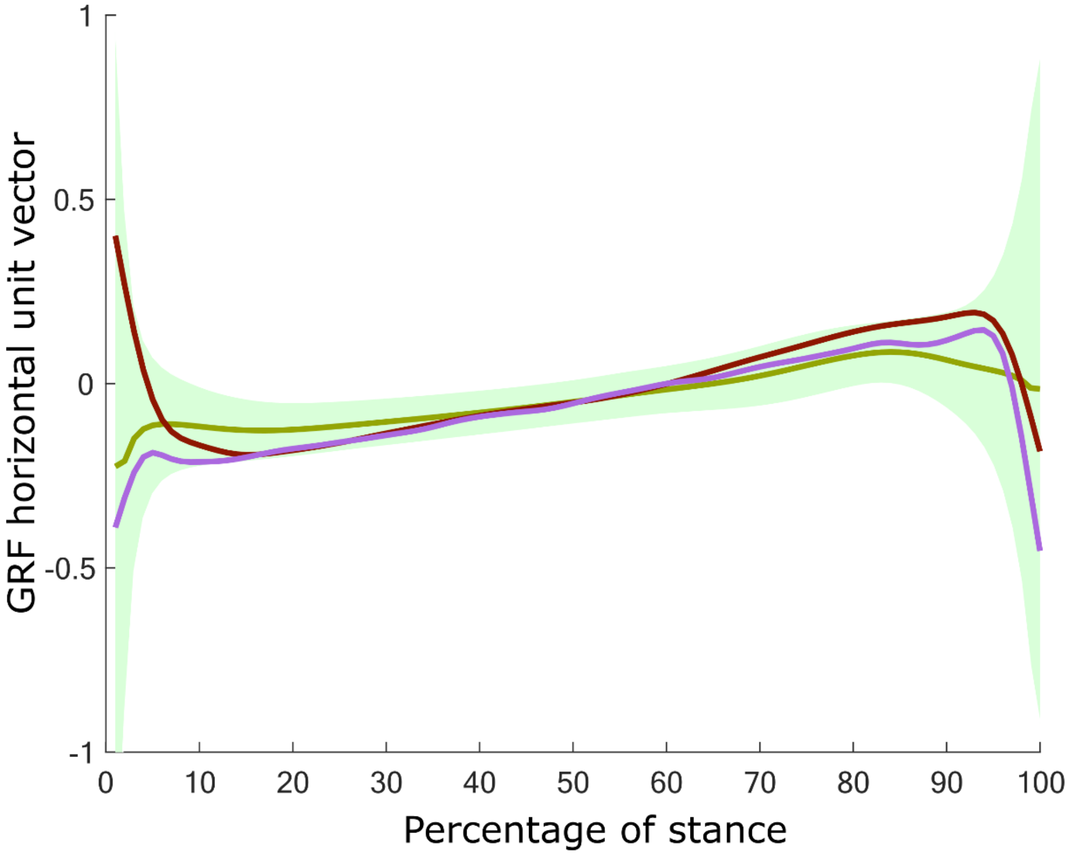


**Figure S7 Horizontal GRF unit vectors during the stance phase, in the giraffe (green), red deer (dark red) and dromedary camel (pink). The mean of 46 trials in the case of the giraffe is shown (1), with the shaded region showing the variation (two standard deviations) observed. The data for the deer and dromedary are from 1 step each (2). The vertical components (not shown) in each species remained consistently >0.97 when the limb was fully loaded, contrasting with the horizontal component which changed sign from a negative (braking) to positive (acceleratory) vector. The unit vectors for red deer and dromedary fell within the variation seen in giraffes.**

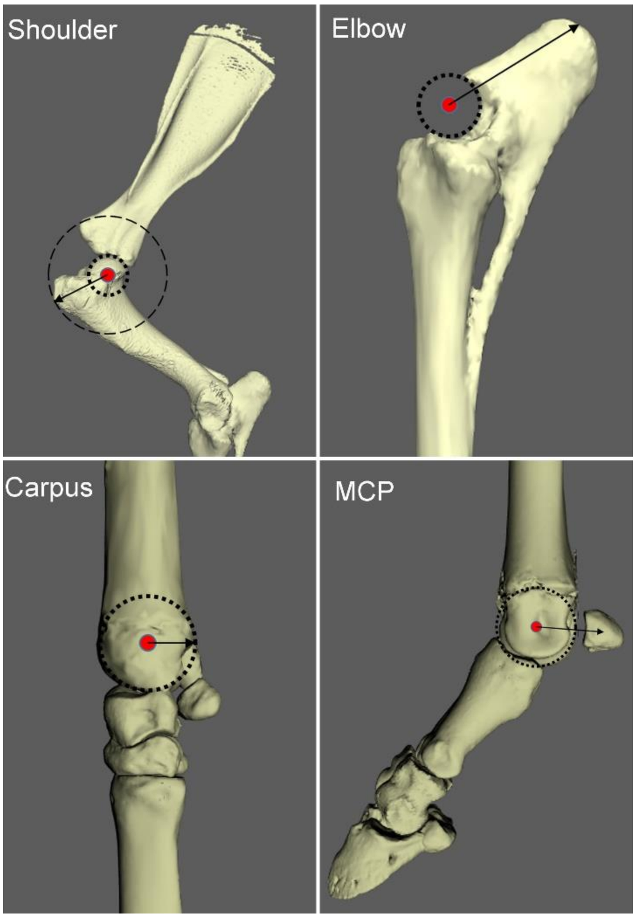


**Figure S8 Bone geometry alone was used as a measure of left forelimb (lateral view) muscle moment arms in *Giraffa* (shown), *Sivatherium* and *Okapia*. Dotted lines represent shape fitting to the joint surface, which facilitated measurement of the moment arms (black arrows) in the parasagittal plane. The shoulder extensor moment arm was defined by the distance from the joint centre to the greater tubercle. This vector is perpendicular to the path taken by the biceps brachii muscle, as it wraps around the shoulder joint. The orientation of the elbow moment arm was defined using a previous methodology (3). The length of the moment arm was defined by the centre point of the corresponding bony region (bounded by the two small red dots). The carpus flexor moment arm was defined by the distance from the centre of rotation to the fitted circle. The MCP flexor moment arm was defined as the distance from the centre of rotation to the middle of the proximal sesamoid bones.**

| Muscle groups active during stance | Total PSCA ($m^2$) | Weighted mean fascicle length (m) | Weighted mean muscle moment arm (m) | Static model muscle moment arms (m) | | |
|---|---|---|---|---|---|---|
| | | | | *Giraffa* | *Sivatherium* | *Okapia* |
| Shoulder extensors | 0.10 | 0.228 | 0.065 | 0.10 | 0.13 | 0.07 |
| Shoulder flexors | 0.076 | 0.114 | 0.14 | Not measured | Not measured | Not measured |
| Elbow extensors | 0.14 | 0.116 | 0.062 | 0.08 | 0.15 | 0.05 |
| Carpal flexors | 0.021 | 0.106 | 0.060 | 0.032 | 0.06 | 0.02 |
| MCP flexors | 0.040 | 0.059 | 0.059 | 0.058 | 0.036 | 0.02 |

**Table S1 Muscle groups with their combined physiological cross sectional area (PCSA), weighted mean fascicle length $l_{fasc}$, and mean muscle moment arm (weighted by each muscle's contribution to PCSA) during the stance phase. Comparable measurements from the static skeletal models are shown for comparison.**

| Specimen number | Description of specimen | Intact/Partial fragment |
|---|---|---|
| NHMUK PV OR 39536 | Left scaphoid carpal (radial carpal) | Intact |
| NHMUK PV OR 39537 | Left lunar carpal (intermediate carpal) | Intact |
| NHMUK PV OR 39538 | Right cuneiform carpal (ulnar carpal) | Intact |
| NHMUK PV OR 15695 | Left magnum carpal (C2+3) | Intact |
| NHMUK PV OR 39540 | Left unciform carpal (C4) | Intact |
| NHMUK PV OR 17089 | Right MC with articulated magnum | Intact |
| NHMUK PV OR 39541 | Proximal phalanx | Intact |
| NHMUK PV OR 15805 | Second phalanx | Intact |
| NHMUK PV OR 39534 | Right radioulnar | Intact |
| NHMUK PV OR 39688 | Left humerus | Intact |
| NHMUK PV OR 36680 | Left scapula | Partial |

**Table S2 Specimen details for *Sivatherium giganteum.* All specimens held at the Natural History Museum, South Kensington, UK.**

| Joint | Internal angle, flexor aspect (°) |
|---|---|
| Shoulder | 105 |
| Elbow | 126 |
| Carpus | 181 |
| MCP | 198 |

**Table S3** ***Okapia johnstoni*** **midstance angles (4)**

### SI References


1. C. Basu, A. M. Wilson, J. R. Hutchinson, The locomotor kinematics and ground reaction forces of walking giraffes. *Journal of Experimental Biology* **222** (2019).
2. S. E. Warner *et al.*, Size-related changes in foot impact mechanics in hoofed mammals. *PloS one* **8**, e54784 (2013).
3. S.-i. Fujiwara, Olecranon orientation as an indicator of elbow joint angle in the stance phase, and estimation of forelimb posture in extinct quadruped animals. *Journal of morphology* **270**, 1107-1121 (2009).
4. K. D'Aout, C. Marien, K. Leus, P. Aerts (2005) Gait patterns and hoof impact in a captive giraffid, the Okapi (*Okapia johnstoni*). in *COMPARATIVE BIOCHEMISTRY AND PHYSIOLOGY A-MOLECULAR & INTEGRATIVE PHYSIOLOGY* (ELSEVIER SCIENCE INC 360 PARK AVE SOUTH, NEW YORK, NY 10010-1710 USA), pp S148-S148.
